# Distinguishing Different Levels of Consciousness using a Novel Network Causal Activity Measure

**DOI:** 10.1101/660043

**Authors:** Nikita Agarwal, Aditi Kathpalia, Nithin Nagaraj

## Abstract

Characterizing consciousness, the inner subjective feeling that is present in every experience, is a hard problem in neuroscience, but has important clinical implications. A leading neuro-scientific approach to understanding consciousness is to measure the complex causal neural interactions in the brain. Elucidating the complex causal interplay between cortical neural interactions and the subsequent network computations is very challenging. In this study, we propose a novel quantitative measure of consciousness - *Network Causal Activity* - using a recently proposed *Compression-Complexity Causality* measure to analyze electrocorticographic signals from the lateral cortex of four monkeys during two states of consciousness (awake and anaesthesia). Our results suggest that Network Causal Activity is consistently higher in the awake state as compared with anaesthesia state for all the monkeys.

## I. Introduction

Understanding *Consciousness* – the inner subjective feeling that is present in every experience (eg., in “seeing” a red rose, in the “feeling” of pain, in the “tasting” of tea etc.), is the final frontier of biomedical research. Defining, modeling and measuring consciousness is considered a *hard* problem [1]. Consciousness largely bounds two facets, namely, the “level” of consciousness and the “content” of consciousness. Experiences such as coma, different stages of anaesthesia and certain stages of sleep seem to indicate a loss of consciousness [2]. Quantitatively, consciousness can be featured as the distributed cortical activity in the sub-cortical regions of the brain relating to the conscious content at any instant. Qualitatively, consciousness is the most essential aspect of our daily experience as it plays a big role in decision making and adaptive planning [3].

Measuring consciousness is a great aid to clinical assessments as it helps in building computational and psychological models; and philosophical aspects to understand the principles connecting brain activity to consciousness experience of wakeful individuals and individuals with physiological, pharmacological and pathological loss of consciousness. Recently, a number of scientific measures of consciousness have been proposed, each having their own theoretical and mathematical framework. We shall briefly describe a few of them here.

Tononi’s Integrated Information Theory of Consciousness (IIT) is a leading scientific theory [4], [5] that conceptualizes the criteria for assessing the consciousness level of any system, quantitatively, as well as, qualitatively. According to IIT, if a system *intrinsically* possesses both *integrated* and *differentiated* states of *information*, then it is bound to possess some level of consciousness (indicated by the symbol Φ in the theory). Non-zero values of Φ conforms the system is in a conscious state. Neurobiologically, the number of connections of neurons in brain networks as well as their complex dynamical interactions contributes to the quantification of Φ, not necessarily only the number of neurons. Perturbational Complexity Index (PCI) [6] is a theory-driven index formulated to evaluate the level of consciousness in a clinical scenario. In order to calculate PCI, the cortex of brain is perturbed with trans-cranial magnetic stimulation to invoke distributed activity in the thalamocortical brain networks. These spatiotemporal responses are then compressed to measure their algorithmic complexity which is normalized and calibrated to yield an index of consciousness level known as PCI. A high value of PCI indicates a high and significant amount of complex interactions of neural activity in cortical areas. Another measure of consciousness, Neural Complexity [7], aims to quantify the interplay between statistically independent (functionally segregated) and statistically interdependent (functionally integrated) neuronal groups in the brain. For any dynamical system, it is an information theoretic measure that captures the mutual information present between the different active subsets of the whole system [8]. Yet another measure of consciousness, known as Causal Density [9], [10], defines consciousness as the fraction of significant causal interactions in brain networks using Granger Causality measure [11]. In [12], [13], there is a review of other scientific measures of consciousness which are similar to the ones described here. The aforementioned measures are categorized under *Complexity Theories of Consciousness* (please see [14]).

In [15], a simple model of spectral Granger bivariate causality is applied to visualize the information flow between different parts of cortex for different states – conscious and unconsciousness induced by different means, in monkeys. This enabled the investigation of large-scale information flow and causal interactions specific to different frequency-modes in the brain. A switch in the frequency-mode of neural communication was found to characterize the difference between different levels of consciousness in monkeys.

In this study, we propose a novel, time domain, *Network Causal Activity* approach to discriminate different levels of consciousness. A recently proposed causality measure, *Compression-Complexity Causality* [16], which has been rigorously tested on simulated data for realistic scenarios as well as real-world data is used to formulate the measure. Instead of frequency domain analysis used in [15], we use this Network Causal Activity measure to differentiate between conscious and unconscious states in monkeys. The organization of the paper is as follows. In section II, we give a description of our proposed methodology, the data sets used, and define Network Causal Activity measure based on Compression-Complexity Causality which is then applied on the data. This is followed by a detailed analysis and discussion in section III. We conclude in section IV with future research possibilities.

## II. Materials and Methods

### A. Subjects and Data Acquisition

For our work, we have used a subset of the dataset from the study conducted by Yanagawa *et al*. [15] which is made available in the public server neurotycho.org (http://neurotycho.org/) [17]. In their study, electrocorticographic (ECoG) signals sampled at 1 kHz were recorded by a Cerebus data acquisition system (Blackrock, UT, USA) from the lateral cortex of four monkeys (George, Chibi, Su, Kin2) using 128 channels electrodes during different stages of sleep, wakefulness and anaesthesia on different days. A complete description about the experiment can be found in [15]. We have focused on only the awake (eyes-opened) and ketamine-medetomidine induced anaesthetized conditions.

### B. Dataset Description

From the recorded neural data collected from the experiments of the study in [15], 3 non-overlapping windows of 5s each (corresponding to 5000 time points) were extracted from 126 channels to construct a sustained network of neural interactions for all the four monkeys in two states – awake (eyes open, conscious state) and anaesthetized (drugged using Ketamine and Medetomidine, loss of consciousness state) condition. We excluded data from two channels (nos. 73 and 123) since these were found to be unsuitable for computation of causality values. The data was used in the acquired form without any re-referencing or pre-processing.

### C. Network Causal Activity

Network Causal Activity (NCA) is proposed as a quantitative measure of consciousness to capture average (significant) causal influence activity between all the elements or subsystems of a given system. We use a recently proposed measure – *Compression-Complexity Causality* (CCC) [16] to estimate causal influences. CCC Toolbox made available as a part of Supplemental Material for [16] was used.

CCC is a time-domain, model-free measure that has been demonstrated to outperform the well-known Granger Causality [11] and Transfer Entropy [18] for several systems (stochastic and chaotic) under wide variety of scenarios, such as, presence of noise, uniform and non-uniform sampling and linear filtering. We shall first briefly describe CCC and then propose NCA using CCC.

For two given time series *A* and *B*, CCC is formulated as follows:

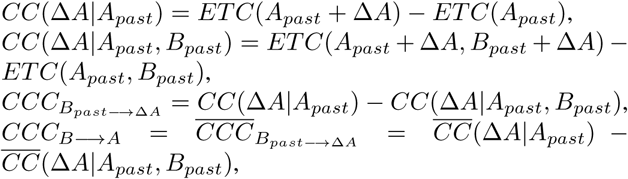

where *CC*(Δ*A*|*A*_*past*_) is the *compression complexity* of time series Δ*A* (the current window with *w* values from time series *A*) given time series *A*_*past*_ (the window of *L* values from the immediate past of Δ*A*). This is estimated by first creating the time series *A*_*past*_ + Δ*A* (here ‘+’ indicates the operation of appending at the end of the time-series) and then computing the difference between the *Effort-To-Compress* the appended time series *A*_*past*_ + Δ*A* and the *Effort-To-Compress* the time series of immediate past values *A*_*past*_. *ETC*(*x*) is a complexity measure that captures the effort required by a lossless compression algorithm (known as Non-Sequential Recursive Pair Substitution Algorithm) to convert *x* to a constant sequence [19]. ETC values are high for random sequences and low for periodic sequences. Similarly, *CC*(Δ*A*|*A*_*past*_, *B*_*past*_) refers to the compression complexity of time series Δ*A* given both the time series *A*_*past*_ and *B*_*past*_ (the immediate past values of Δ*A* taken from time series *A* and *B*). This is computed by taking the difference between *ETC*(*A*_*past*_ + Δ*A, B*_*past*_ + Δ*A*) and *ETC*(*A*_*past*_, *B*_*past*_) where *ETC*(*a, b*) is the joint effort-to-compress complexity of time series *a* and *b. CCC*_*B*_*past*→Δ*A*, which is computed as a difference of two compression complexities (*CC*(Δ*A*|*A*_*past*_) − *CC*(Δ*A*|*A*_*past*_, *B*_*past*_)), is a measure of *causality* from the immediate past of *B* to the current window of *A*. To compute the overall *causality* from time series *B* to *A* we take the average over all the temporal windows yielding *CCC*_*B*→*A*_.

A statistically significant non-zero value of *CCC*_*B*→*A*_ implies a causation from *B* to *A*. A similar estimation of CCC from *A* to *B* can be computed (*CCC*_*A*→*B*_).

Having defined CCC for two time series, Network Causal Activity for multi-variate time series data **M** (with *m* variables^1^) is defined as the total average significant pairwise CCC values across all possible pairs. Mathematically,

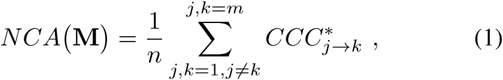

where there are *n* number of *significant* CCC values among all possible pairwise combinations of the *m* variables. CCC value from the *j*-th time series to the *k*-th time series is said to be significant 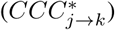 if it is greater than some pre-defined threshold value *T*. Alternatively, we define significance as the highest 10% (in magnitude) of all pairwise CCC values for the given multi-variate time series. These two ways of defining significance will yield different values of NCA. We have used the latter in our study.

We estimated the pairwise CCC values for three non-overlapping windows of ECoG signals of 4 monkeys in *Awake* (conscious) and *Anaesthesia* (loss of consciousness) states. The settings that were chosen for estimating CCC are: *L* = 150, *w* = 30, *δ* = 200 (the step-size for the moving window), *B* = 2 (number of bins^2^). These calculated CCC values are then used to estimate NCA using Eq.1 (*N* = 5000, *m* = 126, *n* = 1575).

## III. Analysis and Discussion

Mean and standard deviation of pairwise CCC values across 126 ECoG signal channels of 4 different monkeys for 3 different windows, each of 5 seconds duration, are given in Table I and Table II for the *awake* and *anaesthesia* states respectively. In Figure 1, histograms of pairwise CCC values for each monkey for window *w*1 (*awake*) and 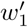 (*anaesthesia*) are shown. In Table III, the Network Causal Activity (NCA) estimates are given for all the monkeys for 3 different windows for both the states. The top 10% significant CCC values were used in computation of NCA. From these tables, we can infer the following:

**Table I:** Mean and standard deviation of pairwise CCC values across 126 ECoG signal channels of different monkeys during awake state for 3 different windows, each of 5 seconds duration. ECoG dataset obtained from [15].

| Monkeys | Awake |  |  |
| --- | --- | --- | --- |
| | CCC: Mean ($\mu$) ± Standard Deviation ($\sigma$) | | |
| | $w_1$ | $w_2$ | $w_3$ |
| George | $0.0165 \pm 0.0176$ | $0.0139 \pm 0.0130$ | $0.0158 \pm 0.0153$ |
| Chibi | $0.0138 \pm 0.0139$ | $0.0159 \pm 0.0156$ | $0.0162 \pm 0.0142$ |
| Su | $0.0183 \pm 0.0164$ | $0.0192 \pm 0.0168$ | $0.0107 \pm 0.0110$ |
| Kin2 | $0.0220 \pm 0.0241$ | $0.0246 \pm 0.0222$ | $0.0186 \pm 0.0196$ |

**Table II:** Mean and standard deviation of pairwise CCC values across 126 ECoG signal channels of different monkeys during anaesthesia state for 3 different windows, each of 5 seconds duration. ECoG dataset obtained from [15].

| Monkeys | Anaesthesia |  |  |
| --- | --- | --- | --- |
| | CCC: Mean ($\mu$) ± Standard Deviation ($\sigma$) | | |
| | $w'_1$ | $w'_2$ | $w'_3$ |
| George | $0.0168 \pm 0.0111$ | $0.0143 \pm 0.0107$ | $0.0140 \pm 0.0106$ |
| Chibi | $0.0073 \pm 0.0074$ | $0.0080 \pm 0.0084$ | $0.0088 \pm 0.0090$ |
| Su | $0.0137 \pm 0.0104$ | $0.0106 \pm 0.0098$ | $0.0140 \pm 0.0116$ |
| Kin2 | $0.0099 \pm 0.0094$ | $0.0114 \pm 0.0099$ | $0.0096 \pm 0.0096$ |

**Table III:** Network Causal Activity (NCA) estimates for all the monkeys for 3 different windows for both awake and anaesthesia states. The top 10% significant CCC values were used in computation of NCA. Mean NCA for awake state is higher than that of anaesthesia.1

| Monkeys | Awake |  |  |  | Anaesthesia |  |  |  |
| --- | --- | --- | --- | --- | --- | --- | --- | --- |
|  | Network Causal Activity |  |  |  | Network Causal Activity |  |  |  |
| | $w_1$ | $w_2$ | $w_3$ | Mean | $w'_1$ | $w'_2$ | $w'_3$ | Mean |
| George | 0.0564 | 0.0427 | 0.0482 | 0.0491 | 0.0378 | 0.0354 | 0.0355 | 0.0362 |
| Chibi | 0.0460 | 0.0512 | 0.0477 | 0.0483 | 0.0214 | 0.0248 | 0.0263 | 0.0242 |
| Su | 0.0555 | 0.0570 | 0.0356 | 0.0494 | 0.0344 | 0.0308 | 0.0391 | 0.0347 |
| Kin2 | 0.0765 | 0.0727 | 0.0632 | 0.0708 | 0.0292 | 0.0307 | 0.0284 | 0.0294 |

**Figure 1:**
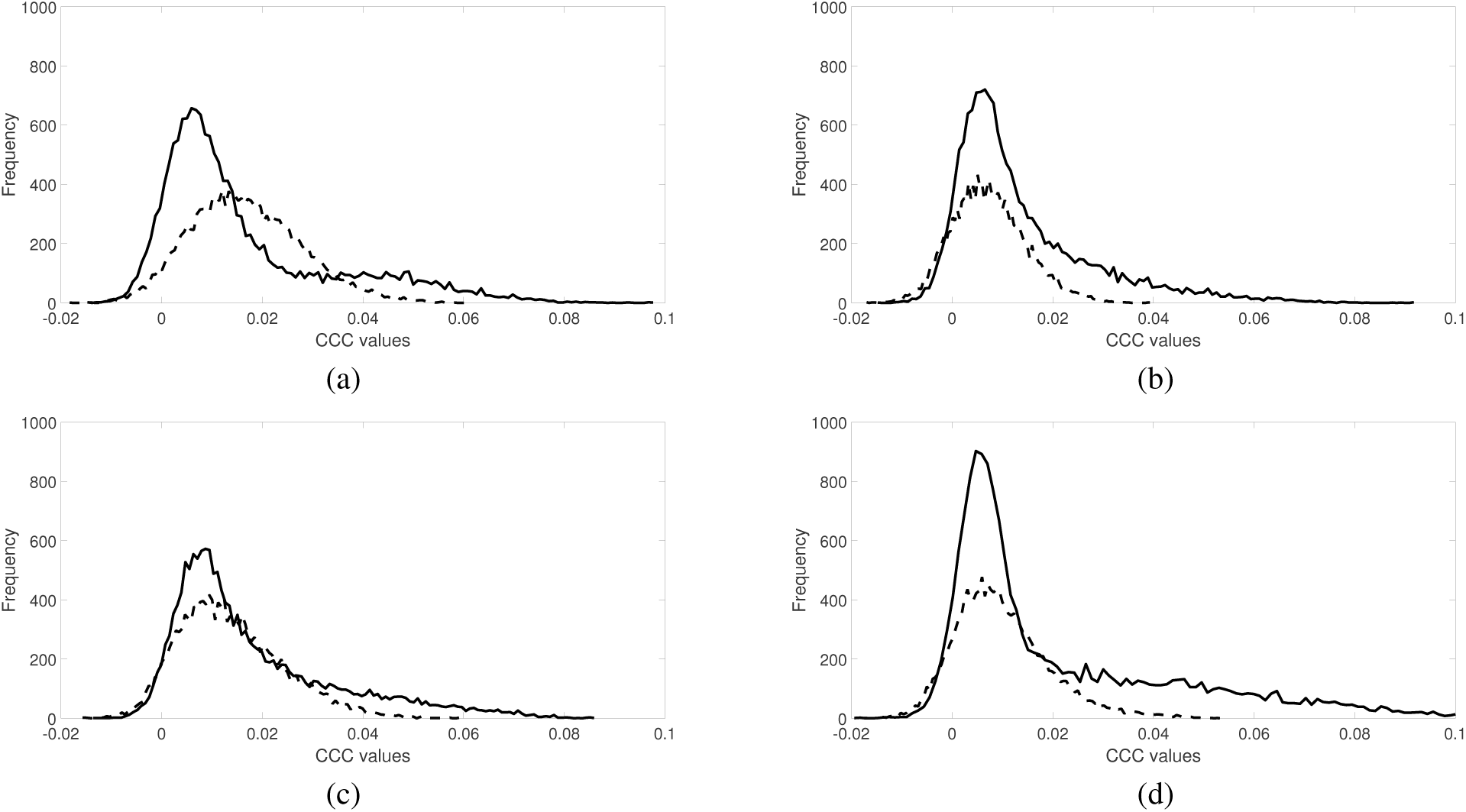
Histogram of pairwise CCC values across 126 ECoG signal channels for each monkey for window *w*_1_ (*awake*) and 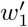 (*anaesthesia*): (a) George, (b) Chibi, (c) Su, (d) Kin2. Solid line (-) is for the *Awake* state and dotted line (--) is for the *Anaesthesia* state. ECoG dataset obtained from [15].

1. It is found that the standard deviations of CCC values in the awake state are *consistently higher* than that of the anaesthesia state in all the windows for all the monkeys (except for one window in case of monkey Su). This finding implies that there are a higher number of differentiated causal neural interactions in the awake state as compared to anaesthesia state.
2. Mean NCA is *consistently higher* in awake state as compared to anaesthesia state across all the windows for all the monkeys. This is intuitive, since in the awake state we expect the average significant causal neural interactions to be of a higher magnitude.
3. The mean CCC values for awake state is significantly higher than the mean CCC value for the anaesthesia state. In order to substantiate this result statistically, a formal hypothesis ‘2 sample student’s t-test’ was performed for all the monkeys on data pooled over all the three windows of awake (*w*_1_, *w*_2_, and *w*_3_) as well as anaesthesia (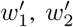, and 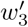). The t-test results are summarized as follows:

- For George, the mean of awake state (0.0154 ± 0.0155) is significantly greater (*t*_94498_ = −4.5272, *p* = 0) than that of anaesthesia state (0.0150 ± 0.0109).
- For Chibi, the mean of awake state (0.0153 ± 0.0146) is significantly greater (*t*_94498_ = −93.7679, *p* = 0) than that of anaesthesia state (0.0081 ± 0.0083).
- For Su, the mean of awake state (0.0161 ± 0.0155) is significantly greater (*t*_94498_ = *-*38.0216, *p* = 0) than that of anaesthesia state (0.0128 ± 0.0107).
- For Kin2, the mean of awake state (0.0217 ± 0.0222) is significantly greater (*t*_94498_ = −103.0372, *p* = 0) than that of anaesthesia state (0.0103 ± 0.0097).

A graphical analysis of this hypothesis test for ‘George’ is depicted in Figure 2.

**Figure 2:**
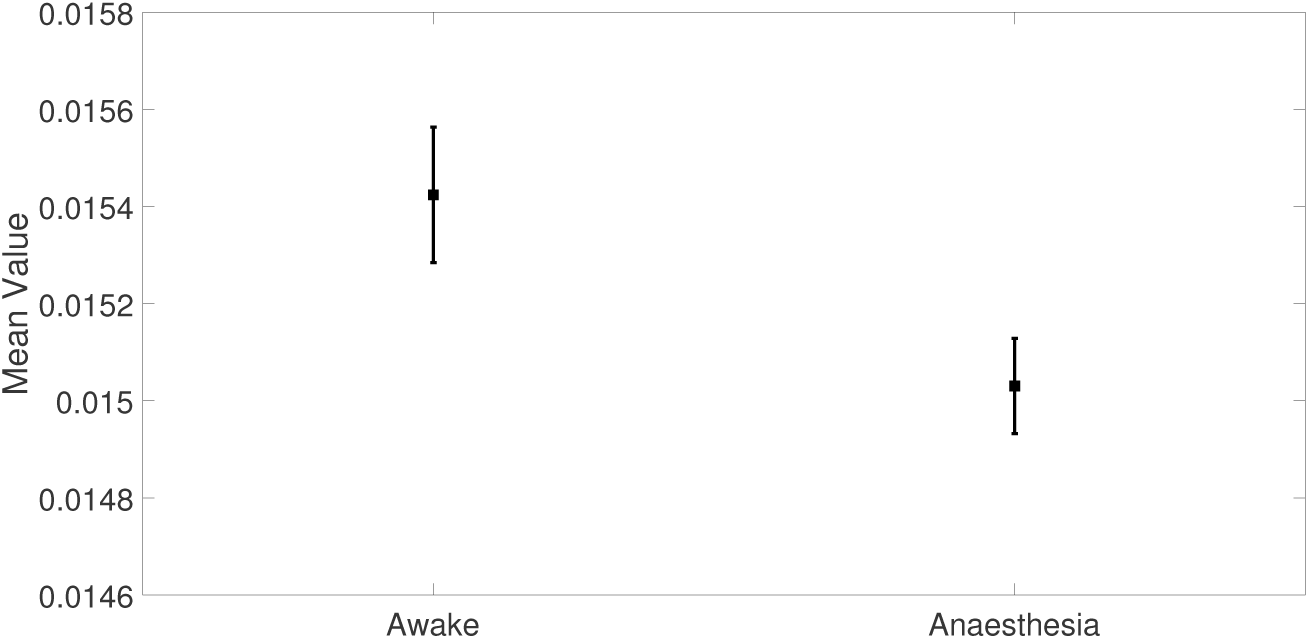
95% confidence intervals for mean CCC values of pooled data of all windows (each of 5 second interval) of George accounting for 47, 250 samples, for *awake* as well as *anaesthesia* state showing a clear separation between the two.

## IV. Conclusion

Measuring Network Causal Activity, i.e., the average significant causal interactions in the brain, is a promising approach towards understanding consciousness. Our work demonstrates that Network Causal Activity, measured by estimating compression-complexity causality values of ECoG signals in monkeys can differentiate states of consciousness (awake vs. anaesthesia). Both, mean CCC and mean NCA measures are statistically significantly higher for awake state when compared with anaesthesia state. Going forward, it is worthwhile to estimate NCA for different stages of sleep and other states of consciousness (such as coma, vegetative state). Potentially, NCA could be further developed to provide robust measures of consciousness in clinical applications.

## Acknowledgment

The authors gratefully acknowledge the financial support of ‘Cognitive Science Research Initiative’ (CSRI-DST) Grant No. DST/CSRI/2017/54, ‘Science and Technology for Yoga and Meditation’ (SATYAM-DST) Grant No. DST/SATYAM/2017/45(G) and Tata Trusts provided for this research. Aditi Kathpalia is thankful to Manipal Academy of Higher Education for permitting this research as part of the PhD programme.

## Footnotes

1 If each of these series has *N* time samples, then **M** is a *m*× *N* matrix. There would be *m*^2^ − *m* pairwise CCC values out of which the highest *n* = 10% are taken as *significant*.

2 CCC is computed on a quantized version of the input real time series. The quantization is performed using uniform sized bins [16].

